# Mechanical activation of TRPV4 channels controls albumin reabsorption by proximal tubule cells

**DOI:** 10.1101/537944

**Authors:** Roberta Gualdani, François Seghers, Xavier Yerna, Olivier Schakman, Nicolas Tajeddine, Younès Achouri, Fadel Tissir, Olivier Devuyst, Philippe Gailly

## Abstract

The proximal tubule (PT) mediates the endocytosis of essential substances filtered through the glomerulus, including albumin and a large variety of low molecular weight proteins that would otherwise be lost in urine. Variations in the amount of ultrafiltrate delivered to the PT modulate protein endocytosis. Here we show that TRPV4 ion channel is expressed at the basolateral side of PT cells. Mechanical activation of TRPV4 by cell stretching induces an entry of Ca^2+^ into the cytosol, which promotes endocytosis. *Trpv4*^−/-^ mice present only a mild PT dysfunction in basal conditions but they exhibit a much more severe proteinuria than *Trpv4*^+/+^ mice when the permeability of glomerular filter is altered by systemic delivery of angiotensin II or antibodies against the glomerular basement membrane. These results emphasize the importance of TRPV4 channel in PT pressure sensing and provide insights into the mechanisms controlling protein reabsorption and potential targets for treating tubular proteinuria.

**Summary:** The proximal tubule (PT) mediates the endocytosis of albumin and low molecular weight proteins. Gualdani et al. report that variations in the amount of ultrafiltrate delivered to the PT activate TRPV4 ion channel expressed at the basolateral side of PT cells, which modulates protein endocytosis.

## Introduction

The proximal tubule (PT) is the first segment of the kidney tubule and it reabsorbs about 70% of the water, ions and solutes filtrated by the glomerulus, using transcellular and paracellular transports. The epithelial cells lining the PT express multiligand receptors megalin (low density lipoprotein-related protein 2, LRP2) and cubilin, that mediate the endocytosis of a large variety of substrates, among which albumin and low molecular weight proteins (LMWP) (Christensen et al., 2009; Devuyst and Luciani, 2015; Eshbach and Weisz, 2017; Raghavan and Weisz, 2016). Albumin is actually reabsorbed via both megalin/cubilin receptor-mediated clathrin-coated pits into vesicles and by fluid-phase endocytosis. The progressive acidification of endosomes can induce a dissociation of albumin from megalin/cubilin receptors and enhance its binding to the neonatal Fc receptor (FcRn). Albumin can thus be sorted either to a megalin/cubilin directed lysosomal degradation or to a FcRn-mediated transcytotic pathway allowing its recycling at the basolateral side and explaining its particular long half-life (19 days in humans) (Kim et al., 2006).

The importance of urinary albumin and LMWP in disease progression is well known, but the mechanisms mediating the presence and toxic effects of albuminuria remain to be determined. The respective importance of the glomerular filtration barrier and the PT function in this process has been recently reexamined. Different studies show that, under physiological conditions, the filtration of albumin is much greater than previously thought and therefore emphasize the role of PT in minimizing albuminuria through reabsorption of albumin (Nielsen and Christensen, 2010; Raghavan and Weisz, 2016).

Variations in the glomerular filtration rate (GFR) have been shown to modulate PT function. Indeed, elevation of GFR increases in Na^+^ reabsorption by inducing the insertion of transporters into the apical membrane (Duan et al., 2010). Increased fluid flow also seems to modulate LMWP endocytosis although this has been shown only on immortalized cell culture models of the PT (reviewed in (Raghavan and Weisz, 2015)). The modulation of PT functions by the amount of ultrafiltrate delivered to the PT suggests the presence of mechanotransduction structures. The ultrafiltrated volume propelled into the PT creates a flow at the apical surface of the PT and raises the intraluminal pressure at a mean of 10-15 mmHg (Drumond and Deen, 1991) but the flow is pulsatile, with an amplitude and a frequency varying according to filtration pressure and heart rate. Two types of force are thus developed by the entry of the ultrafiltrate in the PT lumen: a shear stress on the apical membrane and a radial stretch as changes in intraluminal hydrostatic pressure modify the tubule diameter. Studies performed on immortalized PT cells suggest that the shear stress could be sensed by the primary cilium (Raghavan and Weisz, 2015). Radial stretch for its part is sensed by cytoskeleton, adhesion proteins and/or stretch-sensitive ion channels at the basolateral or apical sides of the epithelial cells (Raghavan and Weisz, 2016). The best candidate channels able to transduce mechanical inputs into ion currents under physiological conditions seem to be Piezo1, Piezo2 and TRPV4 channels (Martinac and Poole, 2018). In PT cells, Piezo1 acts as a stretch-activated cationic channel and Polycystin-2 (PC2) inhibits its activity (Peyronnet et al., 2013). The role of TRPV4 in the PT is so far unknown. TRPV4 is a polymodal ion channel (White et al., 2016) that responds to a number of unrelated stimuli, including temperature, osmotic and chemical stimulations (Ciura et al., 2018; Matsumoto et al., 2018; Shibasaki et al., 2014; Shibasaki et al., 2015; Wissenbach et al., 2000). At the physiological level, TRPV4 plays a role in the mechanotransduction pathways of a variety of cells and tissues, including DRG neurons (Suzuki et al., 2003), osteoblatsts and osteoclatsts (Masuyama et al., 2008; Suzuki et al., 2013), chondrocytes (Servin-Vences et al., 2017) and urothelium (Janssen et al., 2011). In the kidney, which is the organ expressing the most TRPV4 channel, it participates to flow sensing in the collecting duct and MDCK cells (Kottgen et al., 2008; Wu et al., 2007). Recently, we also showed that it constitutes the mechanosensor of the juxtaglomerular apparatus, triggering the inhibition of renin secretion when blood pressure increases (Seghers et al., 2016). In the present paper, we report that TRPV4 is abundantly expressed at the basolateral side of PT cells and that its mechanical activation controls receptor–mediated and fluid phase endocytosis of albumin. In TRPV4^−/-^ mice, this function is altered.

## Results

### TRPV4 is expressed and functional in proximal tubule cells

Immunolocalization demonstrated the presence of TRPV4 in PT cells, mainly at the basolateral and very slightly at the apical membranes (Fig. 1a). Immunofluorescence staining of TRPV4 in PTs showed coincident staining with Na^+^/K^+^-ATPase (Fig. 2a), located within the basolateral membrane, and distinct expression from megalin, a reliable marker of apical side of proximal tubule (Fig. 2b). In order to study the function of TRPV4 channel in the PT, we generated *Trpv4*^−/-^ mice and *in vitro* experiments were performed using mouse primary culture of PT (mPTCs) from *Trpv4*^−/-^ and *Trpv4*^+/+^ mice (Festa et al., 2018). The expression of TRPV4 was confirmed at the mRNA and protein levels in microdissected PT and mPTCs (Fig. 1b, c). In patch-clamp recordings of mPTCs from *Trpv4*^+/+^ mice, application of 100 nM GSK1016790A elicited an outwardly rectifying current, which was completely blocked by the specific TRPV4 antagonist HC067047 (1 µM; Fig. 1d). No current was observed in response to GSK1016790A in mPTCs from *Trpv4*^−/-^ mice (Fig. 1e, f).

**Figure 1.**
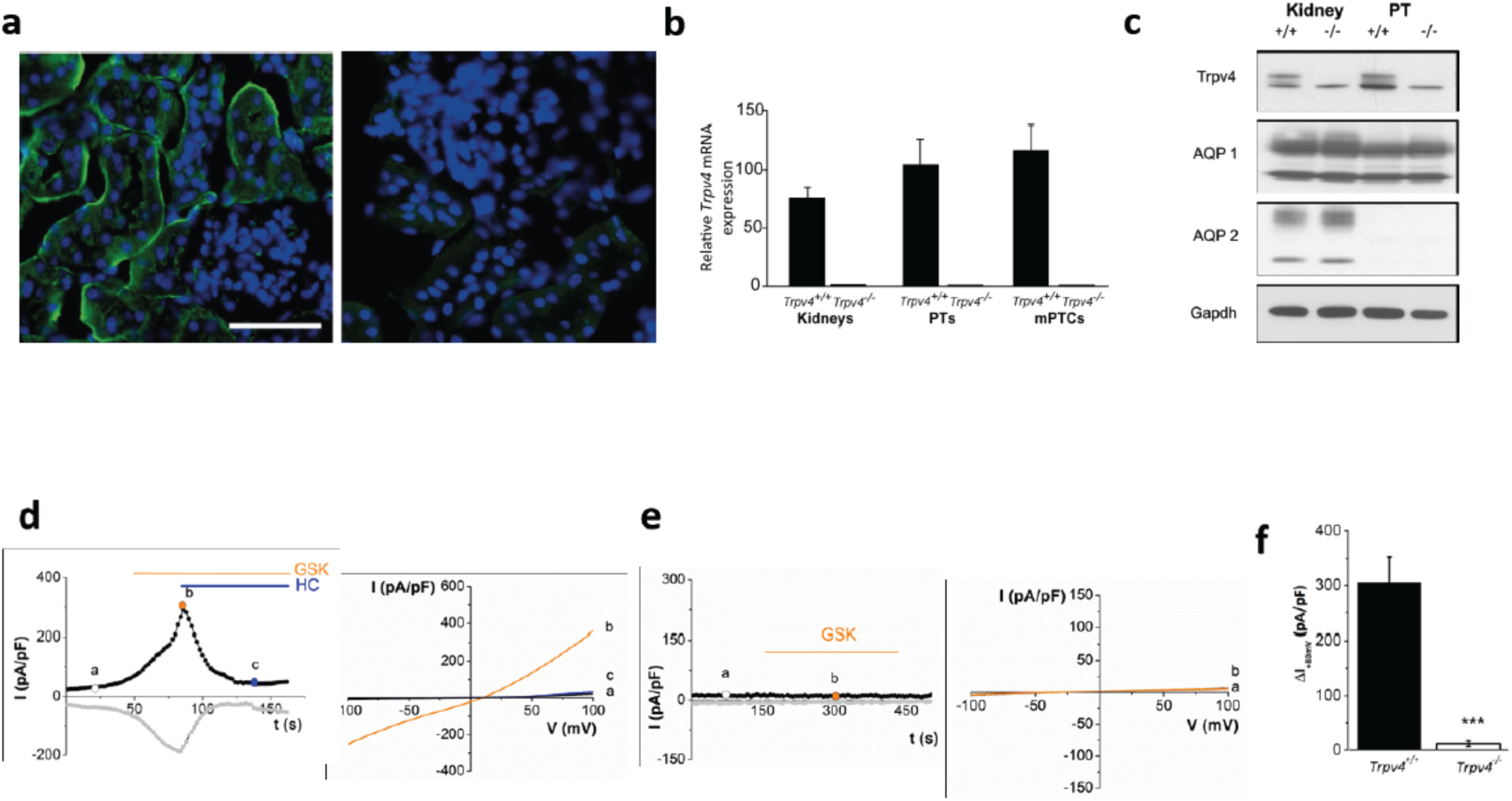
TRPV4 is expressed and is functional in proximal tubule cells. **a)** Immunolabelling of TRPV4 (in green) on renal cortex sections from *Trpv4*^+/+^ (left panel) and *Trpv4*^−/-^ mice (right panel). TRPV4 is mainly expressed at the basolateral side of PT epithelial cells. Nuclei are stained in blue with 1 µg/ml 4′,6-diamidino-2-phenylindole (DAPI). Scale bar: 50 µm. **b)** Quantitative analysis of mRNA expression of TRPV4 channels, in kidneys, proximal tubules (PT) and primary culture from PT (mPTCs), isolated from *Trpv4*^+/+^ and *Trpv4*^−/-^ mice. **c)** Western blotting of total kidney extracts, PT and mPTCs from control (+/+) and knockout (-/-) mice, showing the presence of the TRPV4 protein in the PT and mPTCs of control mice, and its absence in *Trpv4*^−/-^ mice. Anti-AQP1 and anti-AQP2 antibodies are positive and negative markers of proximal tubules, respectively. Anti-GAPDH is used as a loading control. **d)** (left) Representative time-courses of whole-cell current recorded at + 80 mV and −80 mV (black and grey curves, respectively) through *Trpv4*^+/+^ mPTCs, in the presence of 100 nM GSK1016790A and 1 µM HC067047, at the indicated time intervals. (right) I-V traces (from −100 to +100 mV repeated every 2s) obtained at the time points indicated in d). **e)** Same as d), except that mPTCs were isolated from *Trpv4*^−/-^ mice. **f)** Pooled data of whole-cell current (at +80 mV) evoked by 100 nM GSK1016790A, through mPTCs from *Trpv4*^+/+^ and *Trpv4*^−/-^ mice. Each column represents mean ± SEM of n=6 cells. *** p<0.001 (unpaired Student’s t-test).

**Figure 2.**
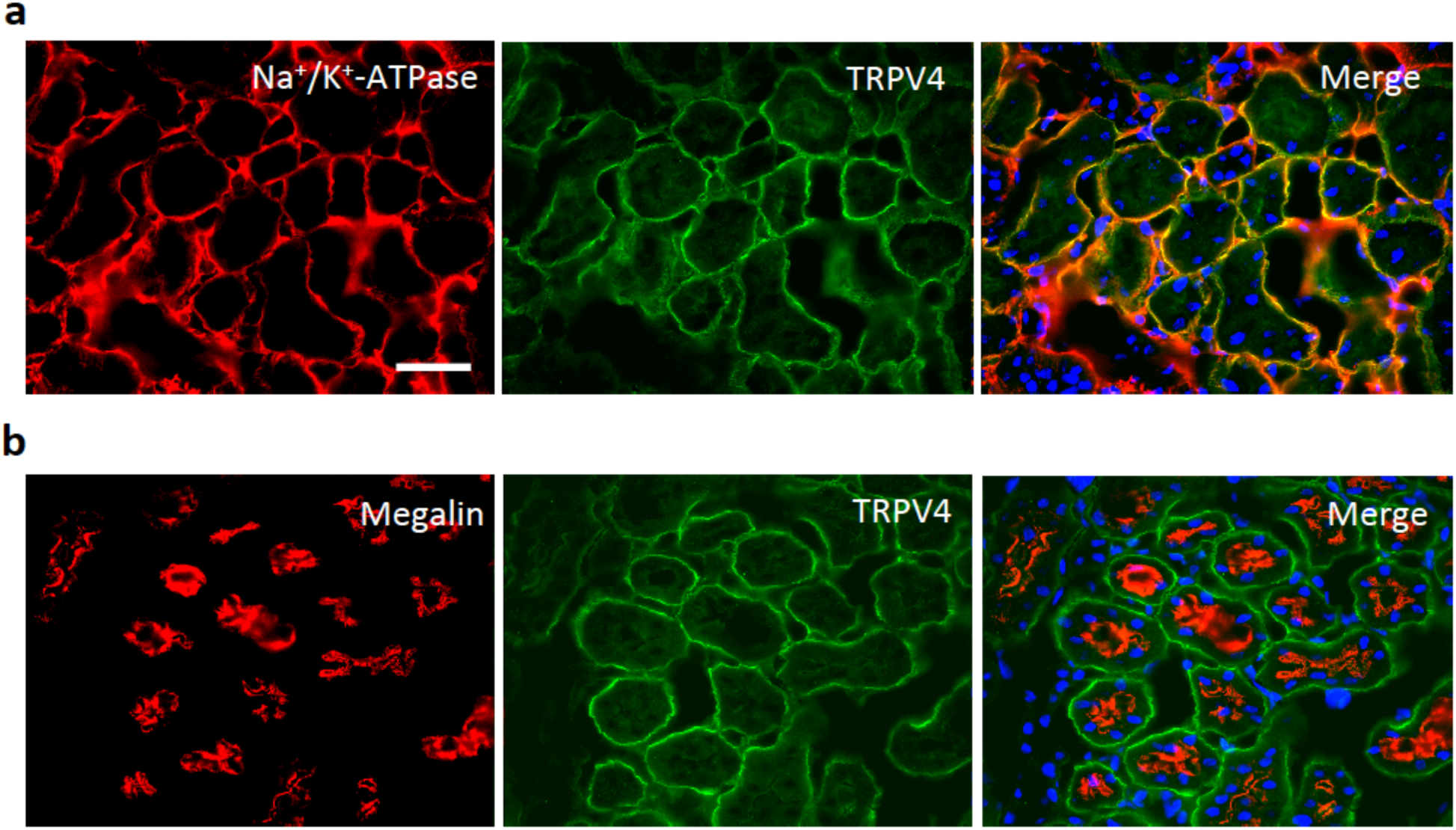
TRPV4 is expressed at the basolateral side of PT cells. Co-immunolabelling of TRPV4 (in green) and Na^+^/K^+^-ATPase **(a**, red) or megalin **(b**, red) on renal cortex sections from *Trpv4*^+/+^ mice. TRPV4 and Na^+^/K^+^-ATPase are mainly expressed at the basolateral side of PT epithelial cells, whereas megalin is expressed at the apical side of PT. Scale bar: 50 µm.

### TRPV4 channel is a mechanosensor in mPTCs

Using Ca^2+^ measurement experiments, we then investigated whether we could mechanically activate TRPV4 in mPTCs. As TRPV4 channel has been described as a flow sensor in the CD (Berrout et al., 2012), we first tested the response of mPTCs to shear stress. When a flow was applied on the surface of the cells, some sparse responses were observed among mPTCs. However, this phenomenon was independent of TRPV4 expression (Fig. 3a, b). We then studied the response of mPTCs to the application of a hypotonic solution. The hypotonic cell swelling induced a [Ca^2+^]_i_ transient in mPTCs that was absent in *Trpv4*^−/-^ cells (Fig. 3c, d). This could indicate that TRPV4 would act as a stretch sensor in the PT. In order to investigate the response of mPTCs to stretch, we cultured them on stretchable silicon chambers. Stretch of the cells induced a rise in [Ca^2+^]_i_. In *Trpv4*^−/-^ cells, the response to stretch presented a significant decrease in the amplitude compared to wild-type cells (Fig. 3e, g). Finally we observed that this response depended on the presence of extracellular Ca^2+^, indicating a Ca^2+^ entry through a mechanically activated channel (Fig. 3f).

**Figure 3.**
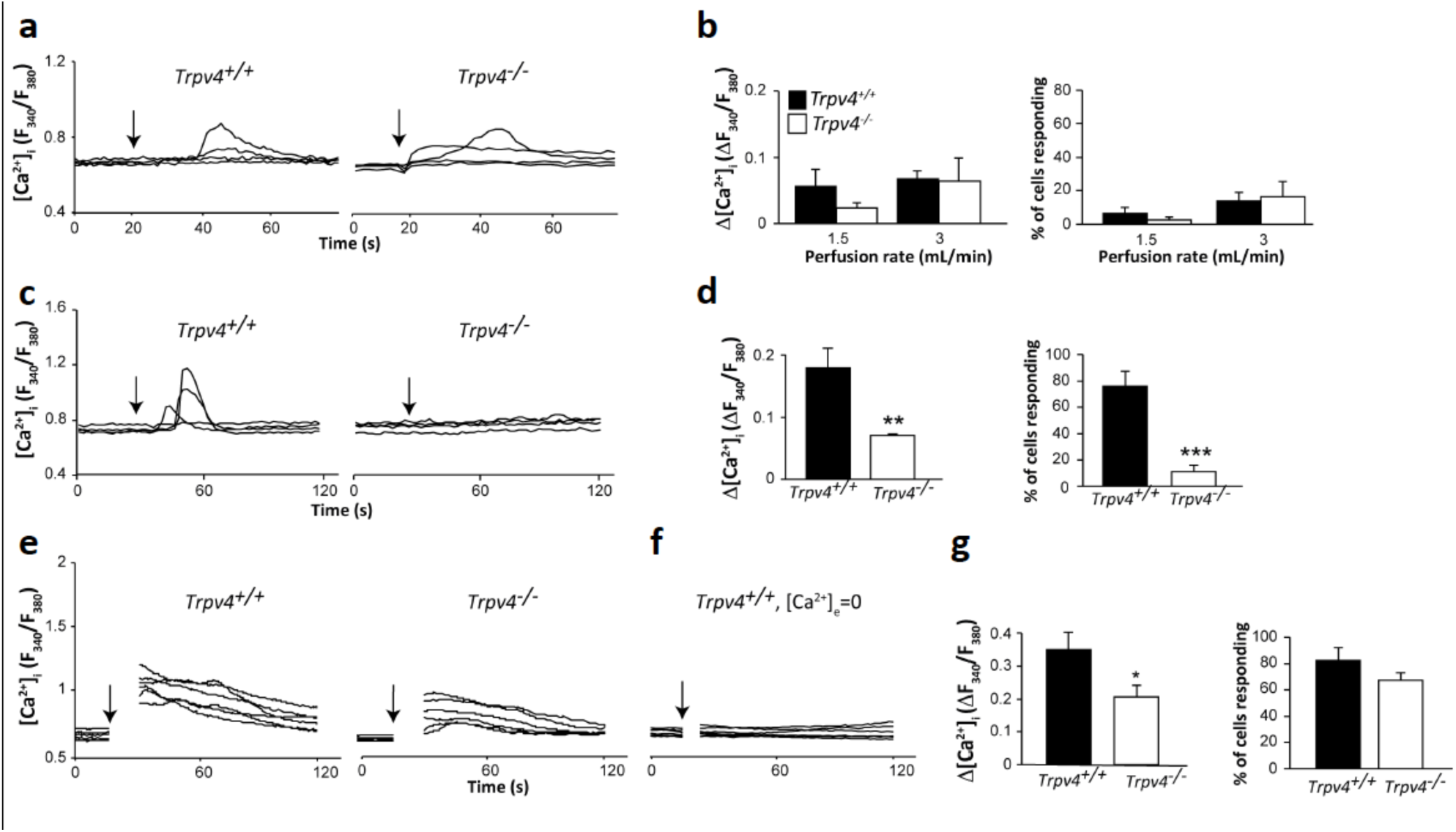
Mechanical response of mPTCs. Changes in [Ca^2+^]_i_ were assessed during flow stimulation **(a, b)**, hypotonic cell swelling **(c, d)** and cell stretching (**e-g)**. For each stimulus, representative curves of individual mPTCs are shown for *Trpv4*^+/+^ and *Trpv4*^−/-^ cells (a, c, e). In f) representative curves of individual *Trpv4*^+/+^ mPTCs stretched in absence of extracellular calcium (*Trpv4*^+/+^, [Ca^2+^]_e_=0) are shown. Average increase in [Ca^2+^]_i_ and proportion of responding cells upon flow stimulation, hypotonic cell swelling and cell stretching (b, d, g) (n=5 experiments with 15-20 cells). * *P*<0.05, ** *P*<0.01, \*\*\**P*<0.001 compared to *Trpv4*^+/+^ cells (unpaired Student’s t-test).

### TRPV4 channel is an osmosensor in mPTCs

TRPV4 was originally identified as a channel activated by hypotonic cell swelling (Liedtke et al., 2000; Strotmann et al., 2000; Vriens et al., 2004; Wissenbach et al., 2000). In *Trpv4*^+/+^ mPTCs, a robust current activation in response to hypotonic shock was recorded by patch-clamp experiments (Fig. 4a). However, a current was also detected in mPTCs derived from *Trpv4*^−/-^ mice (Fig. 4b). This current had the properties originally described for the I_Cl, swell_ (Nilius et al., 1997; Ullrich et al., 2006), i.e. it had a reversal potential close to the predicted Cl^−^ equilibrium potential (∼ −20 mV, Fig. 4g); it was outwardly rectified; it was reverted in the presence of a high concentration of the Cl^−^ channel blocker DIDS (4,4’-Diisothiocyano-2,2’-stilbenedisulfonic acid). Fig. 4a shows that in *Trpv4*^+/+^ mPTCs, the current elicited by hypotonic cell swelling was partially blocked by 100 µM DIDS. On the contrary, the current activated by hypotonic shock in mPTCs from *Trpv4*^−/-^ mice was completely blocked by 100 µM DIDS (Fig. 4b). Interestingly, in *Trpv4*^+/+^ mPTCs, the residual current remaining after application of DIDS showed a reversal potential close to 0 mV, suggesting a different ion selectivity, compared to the unblocked current (Fig. 4a). To test if DIDS impaired the functionality of TRPV4 channel, we perfused *Trpv4*^+/+^ mPTCs with a hypotonic solution containing 100 µM DIDS. This solution activated an outwardly rectifying current, which was completely blocked by the TRPV4 antagonist HC067047 (Fig. 4c). Notably, the reversal potential of this current was ∼0 mV. Of note, the response to 100 nM GSK1016790A was not inhibited by 100 µM DIDS. Our data seemed to indicate that the residual current after pharmacological block of chloride conductance in response to hypotonic shock in *Trpv4*^+/+^ mPTCs was due to the activation of TRPV4 channel. As a matter of fact, a ‘low chloride’ hypotonic solution evoked an outwardly rectifying current, with a reversal potential of ∼ 5 mV (Fig. 4h). This current was completely blocked by HC067047 (Fig. 4d) and was significantly smaller in *Trpv4*^−/-^ mPTCs (Fig. 4e, f), suggesting that TRPV4 contributes to the current rise elicited by hypotonic shock in PT.

**Figure 4.**
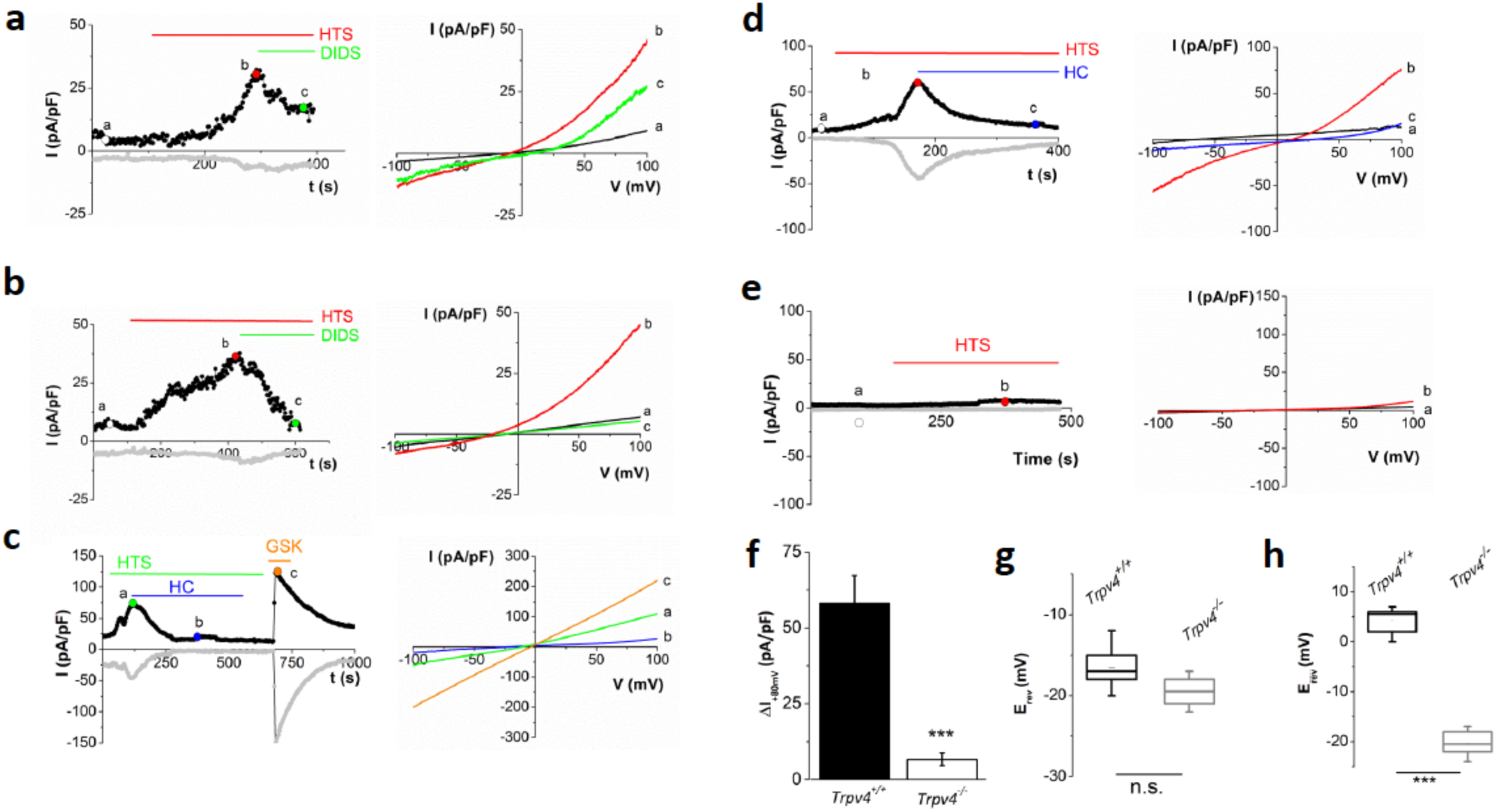
Hypotonic induced-activation of TRPV4 in mPTCs. **a)** Representative time-courses (left) recorded at + 80 mV and −80 mV (black and grey curves, respectively) and I-V traces (right) of whole cell currents through *Trpv4*^+/+^ mPTCs, in the presence of ‘high-chloride’ containing hypotonic solution (HTS) and DIDS (100 µM), at the indicated time intervals. **b)** Same as a), except that time courses and I-V traces are recorded through *Trpv4*^−/-^ mPTCs. **c)** Representative time-courses (left) recorded at +80 mV and −80 mV (black and grey curves, respectively) and I-V traces (right) of whole cell currents through *Trpv4*^+/+^ mPTCs, during administration of hypotonic solution, HC067047 (1 µM) and GSK1016790A (100 nM), at the indicated time intervals, in the continuous presence of DIDS (100 µM). **d)** Representative time-courses (left) and I-V traces (right) of whole-cell current through mPTCs *Trpv4*^+/+^, during administration of ‘low chloride’ hypotonic solution and HC067047 (1 µM), at the indicated time intervals. **e)** Same as d, except that mPTCs were isolated from *Trpv4*^−/-^ mice. **f)** Pooled data of whole-cell current (at +80 mV) evoked by ‘low chloride’ containing hypotonic solution, through mPTCs isolated from *Trpv4*^+/+^ and *Trpv4*^−/-^ mice. Each column represents mean ± SEM of n=6 cells. *** p<0.001 (unpaired Student’s t-test). **g)** Summary of E_rev_ recordings obtained from all IV curves in *Trpv4*^+/+^ and *Trpv4*^−/-^ mPTCs during administration of ‘high chloride’ hypotonic solution (n=6, n.s.; unpaired Student’s t-test). **h)** Similar to g), but for E_rev_ during administration of ‘low chloride’ hypotonic solution (n = 6, p < 0.001; unpaired Student’s t-test).

### Stretch-activated channel (SAC) activity in mPTCs

To investigate the role of TRPV4 channel in the mechanosensitivity of PT we performed high-speed pressure-clamp experiments on mPTCs. Stretch-activated channel activity was elicited by negative suction pulses of 500 ms and patch-clamp recordings were performed in cell-attached configuration at holding potential of −80 mV. As described previously (Peyronnet et al., 2013), currents were characterized by a fast activation, lack of inactivation and a slow deactivation at the end of the pressure pulse (Fig. 5a). Our data showed no difference in amplitude and kinetics of the SAC currents between *Trpv4*^+/+^ and *Trpv4*^−/-^ mPTCs (Fig. 5a and c). Moreover we tested the response of SACs in the presence of 5 µM GsMTx-4, the only drug known to inhibit cationic SACs specifically, without inhibiting other ion-channel families, e.g. voltage-gated potassium and calcium channels (Bowman et al., 2007; Suchyna et al., 2000). SAC currents were completed blocked by extracellular GsMTx-4 (Fig. 5a, bottom, Fig. 5b). Finally we showed that 1 µM HC067047 had no effect SAC currents in *Trpv4^+/+^ cells* (Fig. 5b). These results show that mPTCs do express channels other than TRPV4 that respond to fast membrane stretching obtained with high-speed pressure-clamp. The currents observed are inhibited by GSMTx-4 toxin and are similar to those described by Peyronnet et al. who attributed them to Piezo-1 channels (Peyronnet et al., 2013).

**Figure 5.**
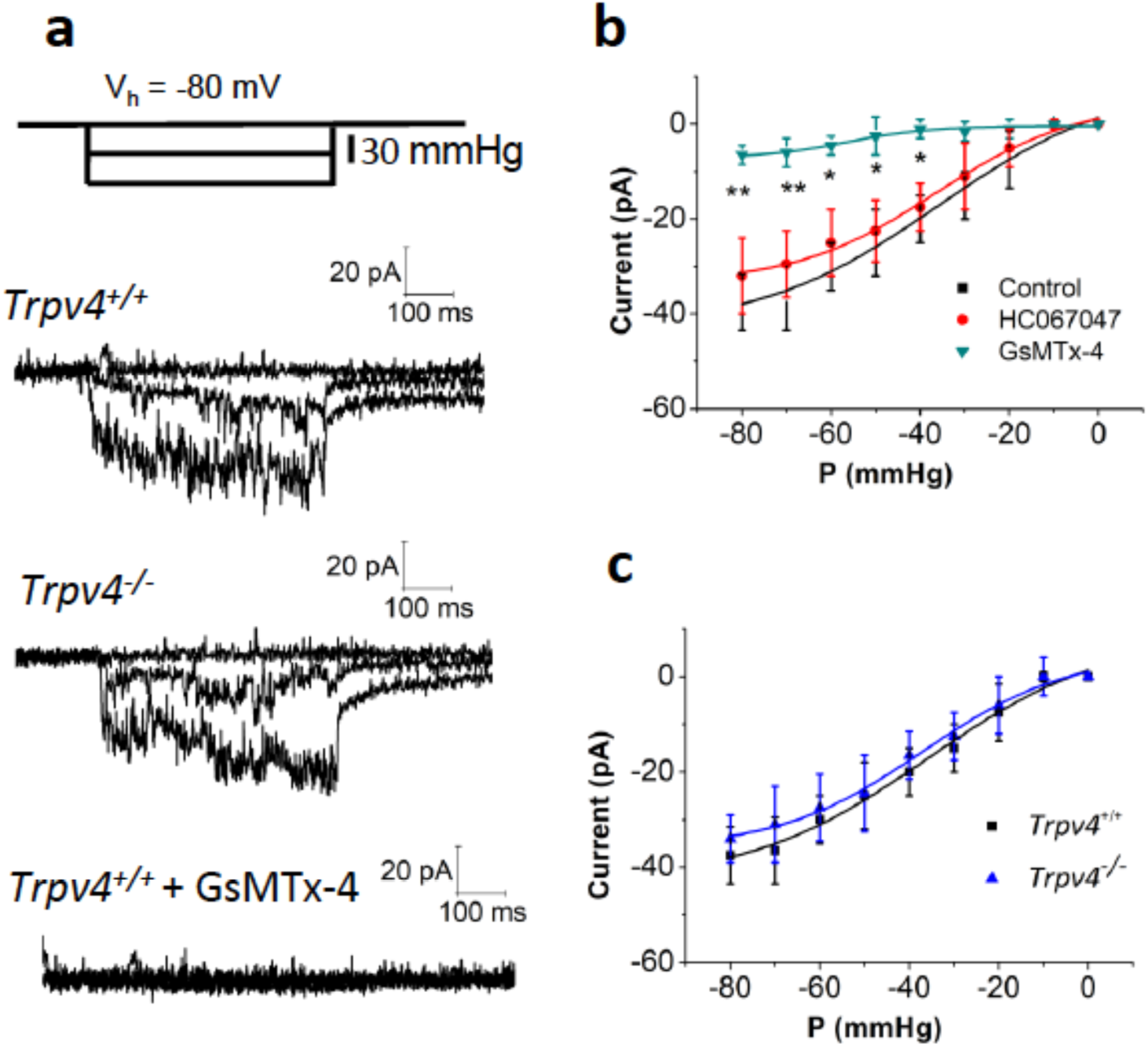
Stretch-activated channel activity in mPTCs. **a)** Cell-attached patch-clamp recording of stretch-activated channel (SACs) activity in mPTCs isolated from *Trpv4*^+/+^ and *Trpv4*^−/-^ mice, in the presence or in the absence of GsMTx-4 (10 µM), at a holding potential of −80mV. Top trace illustrates the pressure pulse protocol. **b)** Normalized current–pressure relationships of SACs in mPTCs from *Trpv4*^+/+^, mice at a holding potential of −80 mV, fitted with a Boltzmann equation. Results are shown as mean ± SEM (n=8 cells). The P_50_ values, extrapolated from the fitting, are: −35,8 ± 4,6 mmHg (control); −36,4 ± 2,7 mmHg (HC067047, 1 µM); −56,4 ± 4,1 mmHg (GsMTx-4, 10 µM) (* *P*<0.05, ** *P*<0.01, ANOVA test). **c)** Same as b), except that SACs are recorded in mPTCs from *Trpv4*^+/+^ and *Trpv4*^−/-^ mice (n=8 cells). The P_50_ value extrapolated from *Trpv4*^−/-^ data is −38,9 ± 3,7 mmHg.

### TRPV4 channel activation triggers albumin and dextran endocytosis in mPTCs

Considering the above results suggesting that PT epithelial cells respond to mechanical cell stretch and the fact that PT functions are controlled by the amount of ultrafiltrate delivered to the PT, we next investigated the possible role of TRPV4 in endocytosis of PT. mPTCs were exposed to FITC-dextran (70 kD) or albumin as markers of fluid phase and receptor-mediated endocytosis, respectively. We quantified the endocytic uptake of albumin and dextran, after pharmacological activation of TRPV4 channel: in the presence of 100 nM GSK1016790A, the uptake of either albumin or dextran was significantly higher in *Trpv4*^+/+^ cells (Fig. 6a-c), compared to the basal level. This effect was not observed in *Trpv4*^−/-^ cells. As expected, the effect of GSK1016790A was abolished in the absence of extracellular Ca^2+^, indicating that Ca^2+^ influx across the plasma membrane is required for increasing endocytosis (Fig. 6d). Notably, stretching the cells increased the albumin uptake, but this effect was blunted in *Trpv4*^−/-^ cells (Fig. 6e). FITC-albumin was highly accumulated in the lysosomes as shown by fluorescence colocalization images of *Trpv4*^+/+^ cells stained with LysoTracker dye after treatment with GSK1016790A (Fig. 6f).

**Figure 6.**
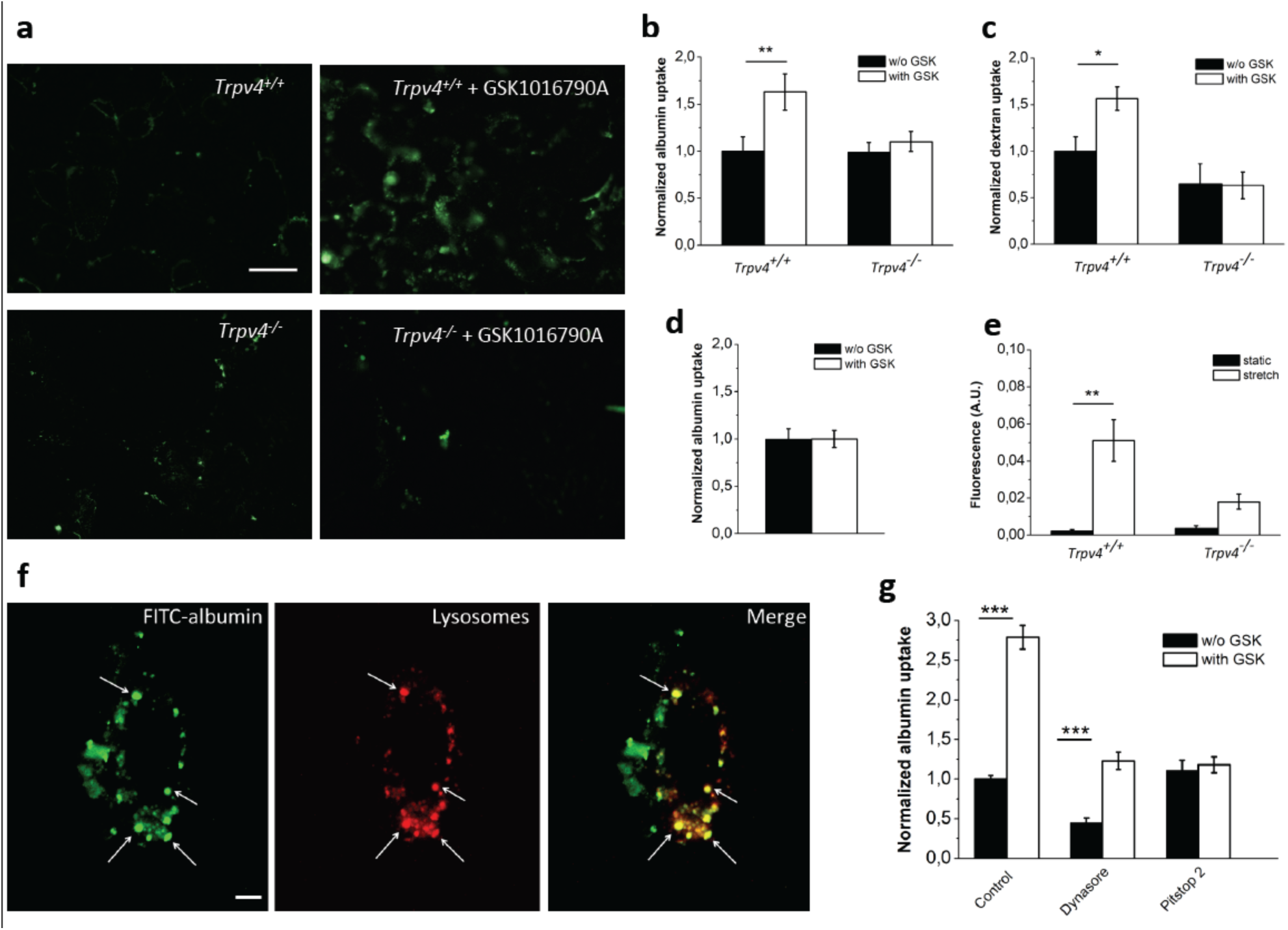
TRPV4 channel determines *in vitro* endocytosis of mPTCs. **a**) FITC-albumin uptake by *Trpv4*^+/+^ and *Trpv4*^−/-^ mPTCs, after incubation with FITC-albumin (0.5 mg/ml) for 15 min., with or without 100 nM GSK1016790A. The scale bar was 50 µm. **b)** Quantification of the total albumin after incubation of mPTCs from *Trpv4*^+/+^ and *Trpv4*^−/-^ mice for 15 min at 37°C, in the absence or the presence of 100 nM GSK1016790A (n= 10, two-way ANOVA with Bonferroni post hoc test ** *P*<0.01). The fluorescence was normalized by the total lysed protein and was expressed as fold-change compared to control. **c)** same as b), except that cells were incubated with 1 mg/ml FITC-dextran for 15 min at 37°C (n= 8, two-way ANOVA with Bonferroni post hoc test * *P*<0.05). **d)** FITC-albumin uptake in mPTCs in Ca^2+^-free medium in the absence or presence of GSK1016790A. **e)** Quantification of albumin uptake in mPTCs from *Trpv4*^+/+^ and *Trpv4*^−/-^ mice incubated for one hour at 37°C with FITC-labelled albumin, in basal and stretched conditions (n=15 fields for each condition from 3 separate experiments; two-way ANOVA with Bonferroni post hoc test * *P*<0.05). **f)** Fluorescence microscopy images showing mPTCs after incubation with FITC-albumin (0.5 mg/ml) and GSK1016790A for 5 min. at 37°C. Lysosomes were stained using LysoTracker Red. The scale bar was 10 µm. **g)** mPTCs were pre-treated with FITC-albumin (0.5 mg/ml) and, where indicated, with 100 µM Dynasore, 30 µM Pitstop-2 or 100 nM GSK1016790A for 30 min. Albumin uptake was quantified as described in Methods, and the mean ± SEM total albumin normalized vs control condition was plotted (n=8, two-way ANOVA with Bonferroni post hoc test *** *P*<0.001).

Reabsorption of albumin from the glomerular filtrate occurs mainly via megalin/cubilin receptor-mediated endocytosis in the PT (Dickson et al., 2014). This process is initiated by binding of albumin to apical clathrin-coated pits. To test if TRPV4-mediated endocytosis in PT occurs via a clathrin- and/or dynamin dependent endocytosis, we preincubated mPTCs with 100 µM Dynasore (dynamin inhibitor) or 30 µM Pitstop (clathrin inhibitor) and, subsequently, added FITC-albumin, in the presence or absence of 100 nM GSK1016790A. Treatment with Dynasore caused a significant inhibition of both basal and TRPV4-induced endocytosis (45% and 44%, respectively). Treatment with Pitstop-2 had no effect on basal uptake of albumin, but reduced TRPV4-induced endocytosis by 55%.

### TRPV4 channel plays a role in tubular proteinuria

In (Janas et al., 2016), working on a mouse model developed by Liedtke and Friedman (Liedtke and Friedman, 2003), we reported that, in basal conditions, no major PT dysfunction was evidenced on the basis of urine analysis in *Trpv4*^−/-^ mice. In particular, we did not observe any abnormal glycosuria, phosphaturia or proteinuria at rest in these mice. In the *Trpv4*^−/-^ mouse model developed here, we confirm these results but interestingly, we found a significantly higher urinary concentration of CC16, a protein secreted by bronchial Clara cells and reabsorbed exclusively by receptor-mediated endocytosis in the early segments of the proximal tubule (Fig. 7a) (Bernard and Lauwerys, 1995). The higher concentration of CC16 observed in *Trpv4*^−/-^ mouse in the absence of a frank albuminuria is suggestive of a mild phenotypic dysfunction of the PT. We hypothesized that TRPV4 could be activated during PT mechanical loading, as this triggers balancing processes increasing its reabsorptive activity. In order to increase the luminal protein delivery to the PT to challenge its endocytic capacity, we used a protocol consisting in continuously delivering angiotensin II (AT2) using osmotic mini-pumps in order to increase the permeability of the glomerular filter (Eckel et al., 2011). We collected urine from *Trpv4*^−/-^ and control mice before and during AT2 administration. Diuresis was significantly increased along AT2 infusion, but similarly between *Trpv4*^−/-^ and control mice (Fig. 8b). The creatinine clearance was similarly altered by AT2 treatment (83.4 ± 7 µl/min, n=10 in *Trpv4*^+/+^ mice and 81 ± 1.4 µl/min, n=10 in *Trpv4*^−/-^ mice), indicating a similar impairment of the glomerular filtration. AT2 increased albuminuria in control mice, but interestingly this increase was much larger in *Trpv4*^−/-^ mice (Fig. 8c). As TRPV4 is not expressed in the glomerulus and as the diuresis and the GFR was affected to the same extent in AT2-treated *Trpv4*^−/-^ and *Trpv4*^+/+^ mice, the specific increase of proteinuria observed in *Trpv4*^−/-^ mice is suggestive of an endocytosis defect in *Trpv4*^−/-^ PT (Raghavan and Weisz, 2016). Moreover, no significant differences in phosphate and glucose urinary levels were found between the two genotypes during the AT2 infusion, suggesting a specific effect on endocytosis (Fig 7). Quantitative analyses of different genes involved in proximal tubule endocytosis confirmed that the higher proteinuria observed in *Trpv4*^−/-^ mice after AT2 injection was not caused by a down-regulation of megalin-cubulin receptor or rabankyrin, KIM-1, Dab-2 and Rab5 factors typically involved in the control of endocytosis (Fig. 8) (Eshbach and Weisz, 2017).

**Figure 7.**
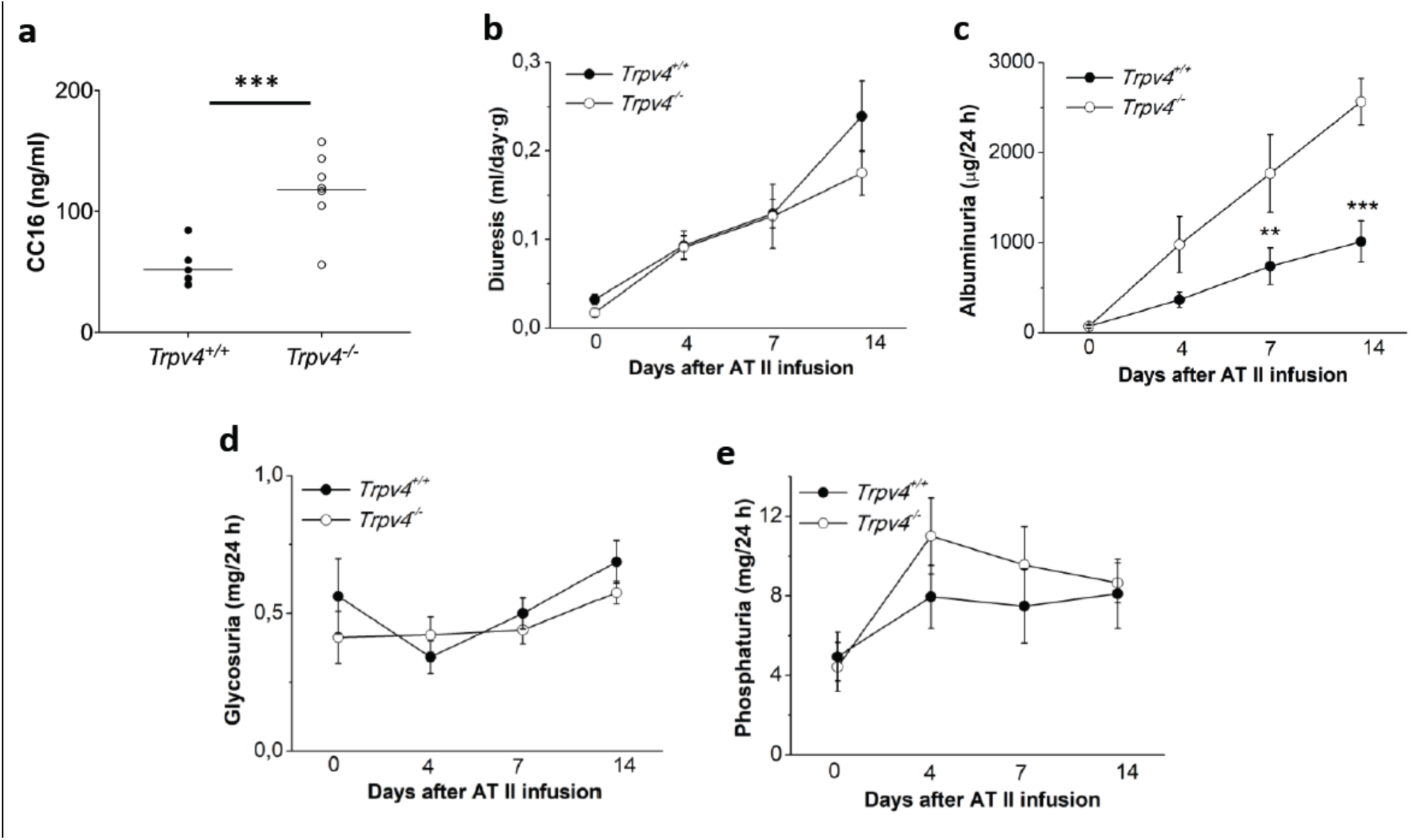
TRPV4 channel participates in *in vivo* albumin-reabsorption after AT2 treatment. **a)** CC16 concentration of basal urines from *Trpv4^−/-^ vs Trpv4^+/+^ mice (P<0,001;* unpaired Student’s t-test). Evolution of diuresis **(b)**, urinary loss of albumin **(c)**, glycosuria **(d)**, phosphaturia **(e)**during AT2 infusion. \*\**P*<0.01 and *** *P*<0.001, *Trpv4^−/-^ vs Trpv4^+/+^*, two-way repeated measures ANOVA (n= 9 pairs).

**Figure 8.**
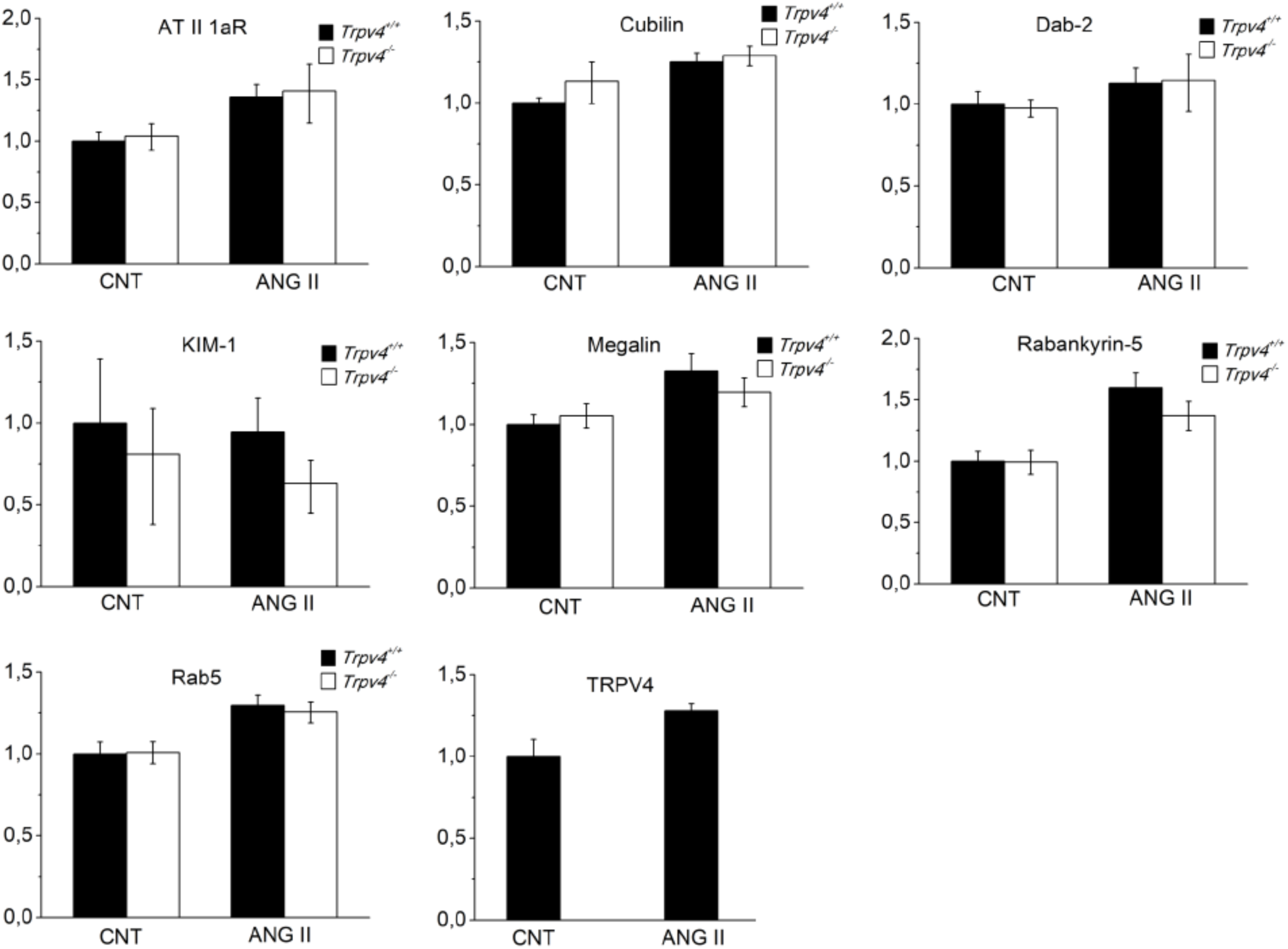
AT2-induced proteinuria is not caused by megalin-cubilin receptor down-regulation. Quantitative analyses of mRNA expression of the following genes, in PTs isolated from *Trpv4*^+/+^ or *Trpv4*^−/-^ mice before or after injection of AT2 using osmotic minipumps: Angiotensin II receptor (AT II 1aR), Cubilin, Disabled-2 (DAB-2), Kidney injury molecule-1 (KIM-1), Megalin, Rabankyrin-5, Rab5, TRPV4.

AT2 infusion has a strong effect on glomerular permeability but it is also associated with many side effects (e.g. hypertension, vascular disease). Therefore, we used a milder and more pathophysiologically relevant protocol to impair glomerular permeability. We proceeded to a unique intravenous injection of 100 µl serum containing antibodies against the glomerular and/or pulmonary basement membrane (anti-GBM serum). Anti-GBM disease is a rare but well-characterized cause of glomerulonephritis, similar to that seen in Goodpasture’s syndrome (Kalluri et al., 1997). It is characterized by the presence of autoantibodies directed at specific antigenic targets within the glomerular and/or pulmonary basement membrane. These antibodies bind to the α3 chain of type IV collagen found in these specialized basement membranes (Sado et al., 1998). Anti-GBM serum treated *Trpv4*^−/-^ mice showed a mild but significant increase of proteinuria, compared to *Trpv4*^+/+^, after 1 week of treatment (*Trpv4*^+/+^: 0.93±0.02 fold change after serum treatment; *Trpv4*^−/-^: 1.43±0.13 fold change after serum treatment. P < 0.05, ANOVA test. Fig. 9). We did not observe such changes in glycosuria and phosphaturia after anti-GBM treatment (Fig. 9b, c).

**Figure 9.**
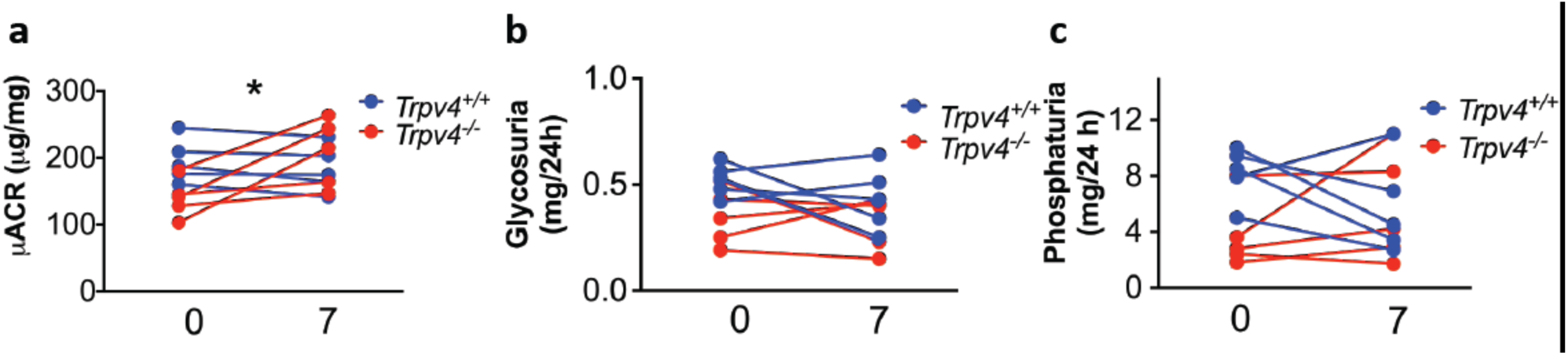
TRPV4 channel plays a role in albumin-reabsorption in a glumerulonephritis mouse model. Evolution of urine albumin-creatinine ratio (µACR) **(a)**, glycosuria **(b)**, phosphaturia **(c)** before and after i.v. injection of anti-GBM serum. Two-way repeated measures ANOVA (n= 5 pairs). Genotype NS, anti-GBM treatment, * *P*<0.05. Interaction ** P<0.01 (increase of proteinuria after anti-GBM treatment only in *Trpv4*^−/-^ mice).

## Discussion

Our data show that TRPV4 channel is expressed in cultured PT cells and can be activated by the specific agonist GSK1016790A and by hypotonic solution. On kidney slices, TRPV4 was immunodetected almost exclusively at the basolateral side of PT cells. As expected, the entry of Ca^2+^ observed in response to shear stress was therefore not affected in mPTCs from *Trpv4*^−/-^ mice. However, a large Ca^2+^ entry was observed in mPTCs in response to mechanical stretch of silicon stretchable chambers and this Ca^2+^ influx is significantly reduced in *Trpv4*^−/-^ PT cells. Surprisingly, in the pressure-clamp experiments, we did not observe any significant difference between *Trpv4*^+/+^ and *Trpv4*^−/-^ mPTCs in response to membrane stretch. In particular, the stretch-activated channel activity in mPTCs was almost completely inhibited by the addition of the toxin GsMTx-4, suggesting that Piezo channels that are sensitive to this toxin, play a critical role in these stretch-activated currents. Interestingly, it was recently shown Piezo1 channel is directly gated by the lipid bilayer tension (Cox et al., 2016), which might not be sufficient for TRPV4 channel. Indeed, at the molecular level, different mechanisms have been proposed for the mechanical gating of TRPV4 channel: i) TRPV4 could be directly activated by membrane stretch, however, it was shown that, in contrast to what is observed *Xenopus laevis* oocytes, TRPV4 overexpressed in mammalian cells do not respond to pressure clamp stimulation (Loukin et al., 2010; Strotmann et al., 2000). ii) The mechanical activation of TRPV4 could be downstream of a PLA2-dependent pathway, as observed after the osmotic stimulus. In this line, shear stress was shown to cause an endothelial dependent vasodilation through PLA2 and TRPV4 activation (Kohler et al., 2006). iii) TRPV4 could be directly activated through deformation of intracellular, transmembrane or extracellular linking proteins. In this context, ultra-rapid activation of TRPV4 (within milliseconds) was observed by magnetically twisting β-integrin in adherence junctions (AJs) (Matthews et al., 2010) and TRPV4 was found in the AJs of the urothelium (Suzuki et al., 2013) and in intercellular junctions between keratinocytes (Sokabe et al., 2010).

Our results support the view that pressure-clamp-induced membrane stretch and uniaxial strain by silicon stretchable chambers stimulate different mechanical pathways in mPTCs and seem in agreement with a recent article that compared the effectiveness of different stimuli on TRPV4 gating in chondrocytes (Servin-Vences et al., 2017). This paper indeed shows that the most effective stimulus to open TRPV4 channel was a membrane deformation at a cell-substrate contact point. The latency was <2ms and the activation was independent of the PLA2 activity, suggesting a direct mechanical gating.

Consistent with (Servin-Vences et al., 2017), we can hypothesize that the strain of silicon chambers is an effective way to gate TRPV4 channel in epithelial PT cells since the stimulus is applied at cell-substrate contact points. However, the identity and the functions of the putative tethers are still to be determined.

Interestingly, the secondary structure of TRPV4 (Nilius and Voets, 2013) and its recently described crystal structure (Deng et al., 2018) show that a large subset of disease-causing mutations are segregated at the outer perimeter of the tetrameric channel, suggesting that the interactions between TRPV4 and protein partners or signaling molecules are crucial for the physiological function of TRPV4, and their alteration could be one of the factor affecting a large number disease phenotypes. The modulation of PT functions by the amount of ultrafiltrate delivered to the PT suggests the presence of mechanotransduction structures. Two types of force are theoretically developed by the entry of the ultrafiltrate in the PT lumen, a shear stress on the apical membrane and a radial stretch on basolateral or apical sides of the PT cells (Drumond and Deen, 1991; Du et al., 2004; Raghavan and Weisz, 2016).

It is known that an increase in glomerular filtration lead to increased fluid shear stress on PT cells that enables the reabsorption of Na^+^ and HCO_3_^−^ ions (Du et al., 2006) and the expression of AQP1 (Pohl et al., 2015). Moreover, a recent study suggests that shear stress enhances endocytosis in PT cells in a Ca^2+^- and cilium-dependent way (Raghavan et al., 2014; Raghavan and Weisz, 2015). In a recent article, however, Delling et al. (Delling et al., 2016) showed that primary cilia from kidney are not Ca^2+^-sensitive mechanosensors. For this reason, further advance in understanding the relationship between Ca^2+^ signal and mechanical modulation of the endocytosis is still needed. In isolated perfused proximal tubules and in unilateral obstruction disease models, increased flow in the PT induces a change in the outer diameter of the tubule as well as the lumen. However, it is so far unclear whether such changes appear under physiological flow variations (Raghavan and Weisz, 2016). Mechanical stretch of PT cells triggers a β-integrin-dependent reinforcement of focal adhesions and an increase of [Ca^2+^]_i_ (reviewed in (Quinlan et al., 2008)). The presence of stretch-activated channels on PTCs has been suggested by Alexander and colleagues who showed that cyclic stretch of isolated PTCs activated PLA2 and extracellular signal-regulated kinases via a mechanism that depends on external Ca^2+^ and is blocked by Gd^3+^ (Alexander et al., 2004). Here, we observed that TRPV4 channel is mainly expressed at the basolateral side of PT. Importantly, we showed that the uptake of FITC-labeled albumin is increased when TRPV4 is activated pharmacologically or mechanically, that the effect on endocytosis depends on the entry of Ca^2+^ in PT cell and that TRPV4-mediated endocytosis occurs via a clathrin- and dynamin-dependent pathway.

We show that *Trpv4*^−/-^ mice present a higher urinary concentration of CC16 a very sensitive marker of PT dysfunction (Bernard and Lauwerys, 1995). Moreover, mice presenting an alteration of the glomerular filter (mouse model subjected to continuous delivery of AT2 or to acute IV injection of anti-GBM serum) develop an albuminuria that is much more important in *Trpv4*^−/-^ than in WT. This suggests the importance of the process in diseases accompanied by altered permeability characteristics of the glomerular filter, some being very rare such as the Goodpasture’s syndrome (see above), some being very common such as the diabetic nephropathy (Lewis and Xu, 2008). PT dysfunction is involved in various forms of Fanconi syndrome, which is often associated with rare inherited diseases such as Dent’s disease, Lowe’s syndrome, Imerslund–Gräsbeck syndrome, cystinosis or Fanconi-Bickel syndrome (Ludwig and Sethi, 2011; Sun et al., 2017). It is characterized by low molecular weight proteinuria, glycosuria, aminoaciduria and / or loss of electrolytes, bicarbonate and lactate. However, PT dysfunction is also largely involved in the proteinuria observed in more common diseases primarily affecting the glomerulus such the early diabetic nephropathy (Gibb et al., 1989; Zeni et al., 2017). Renal tubules are themselves vulnerable to proteinuria and in response to injury, tubular epithelial cells undergo changes and function as inflammatory and fibrogenic cells. This process thus participates to the progression of chronic kidney disease (Liu et al., 2018). The process is also certainly relevant in other disease states accompanied by intraluminal pressure increases, e.g. obstructive uropathy (Cachat et al., 2003; Raghavan et al., 2014; Rohatgi and Flores, 2010; Wyker et al., 1981), polycystic kidney disease (PKD)(Patel and Honore, 2010), renal cysts (Derezic and Cecuk, 1982; Tanner et al., 1995).

We therefore hope that our finding that radial stretch of the PT is sensed via basolateral TRPV4 channels that modify albumin reabsorption will help to treat patients with proteinuria. Future studies will be needed to investigate whether a deficit in TRPV4-mediated endocytosis contributes to kidney diseases with tubular proteinuria.

## Methods

### Mice

*Trpv4*^−/-^ mice were generated using AK7 mouse embryonic stem cells. The *Trpv4* gene was targeted by homologous recombination with a construct from the International KO Mice Consortium that bears *loxP* sites-flanked sixth exon (ID:79331) (Matthews et al., 2010). After selection of several correctly targeted clones, we injected them into C57BL/6J mice host blastocysts. The embryos were transferred into pseudopregnant CD1 mice. In the offspring, chimeric males were selected on basis of the coat color. They were mated with C57BL/6J females in order to assess the transmission of the targeted allele. Mice harbouring the targeted allele were mated with *ROSA*-*Flp* females to have the selection cassette excised. The so obtained mice have the sixth exon of the *Trpv4* gene flanked with *loxP* sites (*Trpv4lox*). *Trpv4^lox/lox^* mice were crossed with a mouse line expressing the Cre recombinase under the PGK-1 promoter to obtain a constitutive *Trpv4* knockout mouse line (*Trpv4*^−/-^). Loss of expression of TRPV4 in *Trpv4*^−/-^ mice was assessed by quantitative RT-qPCR, by Western blotting on kidney or PT extracts and by immunohistochemistry (Fig. 1 a, b and c).

### Ethical approval

All animals were housed and handled according to the Belgian Council on Animal Care guidelines based on protocols approved by the Animal Ethics Committee of the Université catholique de Louvain. Animals were given access to food and water *ad libitum* unless otherwise stated. At appropriate experimental time points, all animals were humanely killed by an overdose of anaesthetic followed by decapitation.

### Immunostaining

*Trpv4*^+/+^ and *Trpv4*^−/-^ mice were deeply anesthetized by IP injection (10 ml/kg) of a solution containing 10 mg/ml ketamine and 1 mg/ml xylazine. After intracardiac perfusion with a 4°C buffered 4% paraformaldehyde PBS solution, the kidneys were removed and postfixed in the same solution for 2 hours, washed in PBS and cryoprotected overnight in 30% sucrose PBS. After embedding in Optimal Cutting Temperature compound (OCT, Tissue-Tek, VWR), 10 µm thick cryosections were cut and stored at −80°C. Sections were blocked with a 3% BSA containing PBS solution and then incubated overnight with 1/200 anti-TRPV4 (Alomone) antibody. Binding sites were revealed with Alexa Fluor 488 (Thermofisher). Images were acquired with an Olympus FV1000 confocal microscope.

### Real-Time PCR

Total RNAs from whole kidney, proximal tubules and proximal tubular cells was extracted using Trizol reagent (Invitrogen) according to the manufacturer’s protocol, and reverse-transcribed using qScript Reverse Transcriptase (Quanta Biosciences, Gaithersburg, ME). cDNA-specific PCR primers were designed using Primer-BLAST (see Table 1) and purchased from Eurogentec (Seraing, Belgium). Gapdh was used as reference gene. Quantitative RT-qPCR was performed using SYBRGreen Mix (Bio-Rad, Hercules, CA, USA). The reaction was initiated at 95 °C for 3 min, followed by 40 cycles of denaturation at 95 °C for 10 s, annealing at 60 °C for 1 min, and extension at 72 °C for 10 s. Data were recorded on a MyiQ qPCR detection system (Bio-Rad), and cycle threshold (*Ct*) values for each reaction were determined using analytical software from the same manufacturer. Each cDNA was amplified in duplicate, and *Ct* values were averaged for each duplicate. The average *Ct* value for *Gapdh*, selected as reference gene for kidney, was subtracted from the average *Ct* value for the gene of interest. The *ΔCt* value obtained in siRNA condition was then subtracted from the *ΔCt* value obtained in control condition, giving a *ΔΔCt* value. As amplification efficiencies of primer pairs were comparable, the *Gapdh*-normalized expression of mRNAs was given by the relation 2^−ΔΔCt^.

**Table 1.**
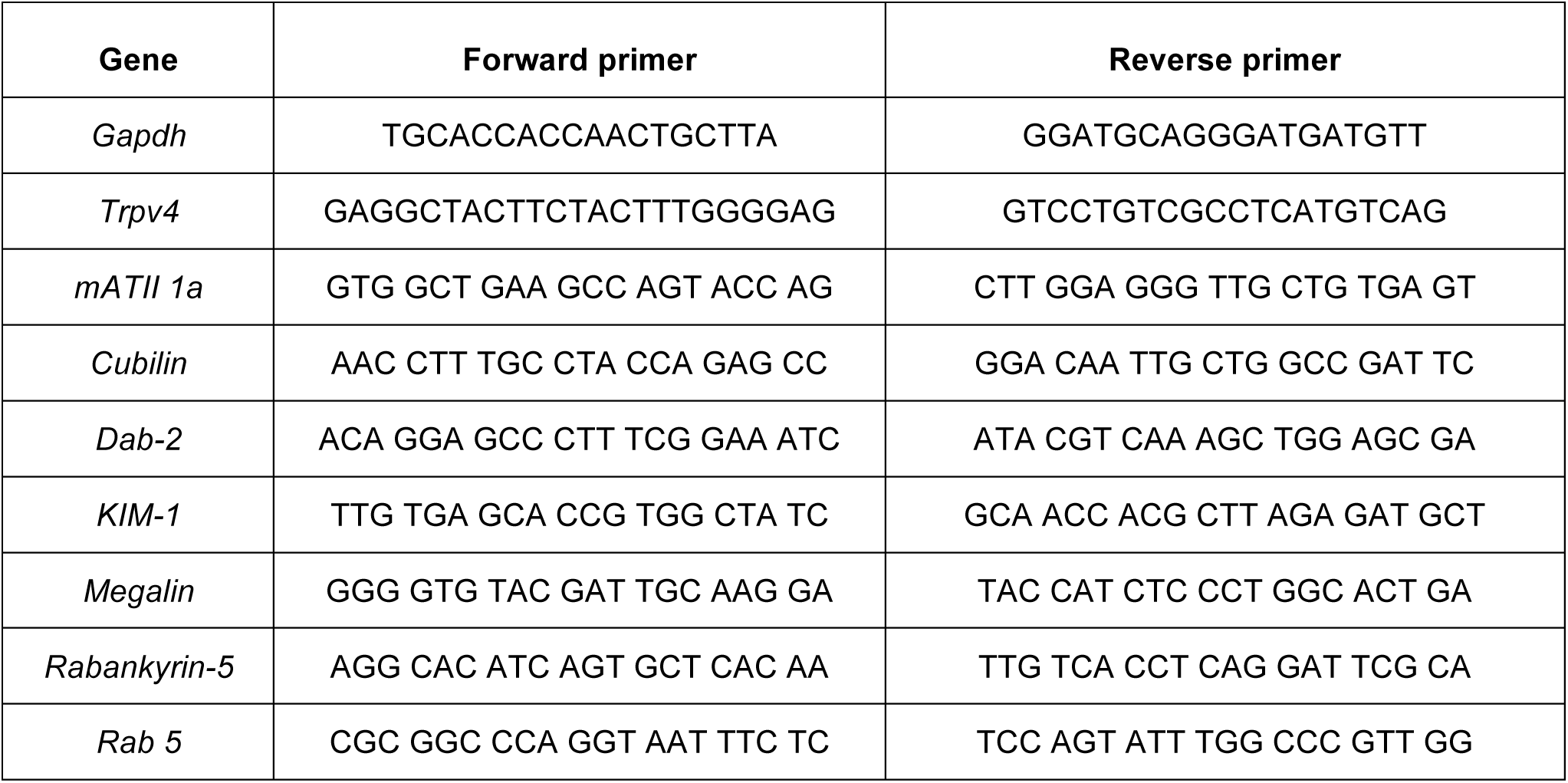
List of DNA primers used in quantitative RT-PCR.

### Western blotting

Cells were collected in RIPA buffer (25 mM Tris HCl pH 7.6, 150 mM NaCl, 1% NP-40, 1% sodium deoxycholate, 0.1% SDS). 30 µg of protein for each sample was loaded on a 10% SDS-polyacrylamide gel, transferred to a PVDF membrane and probed with the following antibodies: a homemade anti-TRPV4 antibody described earlier (Gevaert et al., 2007), an anti-aquaporin 1 antibody (Sigma) as positive marker of proximal tubule, an anti-Aquaporin 2 antibody (Sigma) as negative marker of PT (Janas et al., 2016), an anti-megalin antibody (homemade) as apical side marker of PT, an anti-Na^+^/K^+^-ATPase (Sigma) as a basolateral side marker of PT and an anti-GAPDH antibody as loading control (Cell Signaling).

### Primary culture of mPTCs

Male *Trpv4*^+/+^ and *Trpv4*^−/-^ mice were decapitated, kidneys were removed and decapsulated. The dissection was done as described previously (Janas et al., 2016; Terryn et al., 2007). Renal cortices were dissected visually in HBSS and sliced into pieces of ∼1 mm wide. The fragments were transferred to collagenase solution (HBSS with 0.1% type-2 collagenase and 96 µg/ml soybean trypsin inhibitor) at 37°C and digested for 30 min. After digestion, the supernatant was filtered through two nylon sieves (pore size 250 µm and 80 µm) to obtain a large number of long proximal tubule fragments, without substantial contamination of other nephron segments or glomeruli. The longer PT fragments remained in the 80-µm sieve were resuspended with warm HBSS (37°C) containing BSA 1% (wt/vol). The PTs present in the BSA solution were centrifuged for 5 min at 170 g, washed, and then resuspended into the appropriate amount of culture medium: 1:1 DMEM/F12 without phenol red and supplemented with heat-inactivated 1% FCS, 15 mM HEPES, 2 mM L-glutamine, 50 nM hydrocortisone, 5 µg/ml insulin, 5 µg/ml transferrin, 50 nM selenium, 0.55 mM sodium pyruvate, 10 ml/l 100X non-essential amino acids, 100 IU/ml penicillin and 100 µg/ml streptomycin. The solution was buffered to pH 7.4 and adjusted to an osmolality of 325 mOsm. The PT fragments were seeded onto collagen-coated coverslips and left unstirred for 48 h in a standard humidified incubator. The medium was then replaced every 2 days. After 7 days, cell cultures were organized as a confluent monolayer. No differences in cell shape and size or in proliferation were observed between *Trpv4*^−/-^ and *Trpv4*^+/+^ mPTCs.

### Ca^2+^ imaging

Loading of the mPTCs with Fura2-AM was achieved at room temperature in Krebs medium with 5 µM Fura2-AM and 0.01% pluronic acid for 1h. [Ca^2+^]_i_ was measured in 15-20 individual cells using alternative excitation of Fura2-AM (0.5 Hz) at 340 and 380 nm using a Lambda DG-4 Ultra High Speed Wavelength Switcher (Sutter Instrument, Novato, CA, USA). Images were acquired with a Zeiss Axiocam camera coupled to a 510-nm emission filter and analyzed with Axiovision software. Cytosolic free Ca^2+^ concentration was evaluated from the ratio of fluorescence emission intensities excited at the two wavelengths using the Grynkiewicz equation (Grynkiewicz et al., 1985). Ca^2+^ imaging experiments were performed in Krebs-HEPES solution (NaCl 135 mM, KCl 5.9 mM, MgCl_2_ 1.2 mM, CaCl_2_ 1.8 mM, HEPES 11.6 mM, glucose 11.5 mM, pH 7.35). For hypotonic shock experiments, cells were bathed in a modified Krebs-HEPES solution where NaCl was partially omitted in the hypotonic solution (170 mOsm), or replaced by mannitol in the isotonic solution (310 mOsm), in order to avoid changes in ionic gradients. For shear stress experiments, flow was applied on cells with an 18G needle connected to a peristaltic pump perfusion system and placed above the field. For measurement of [Ca^2+^]_i_ during cell stretching, cells were passaged on collagen I-coated flexible silicon chambers (ST-CH-04, B-Bridge), placed on the microscope-mountable stretching instrument (ST-150, B-Bridge) and strained for 1 s by 20% uniaxial stretching (Seghers et al., 2016).

### Electrophysiology recordings

Patch-clamp recordings of mPTCs were carried out at room temperature, using a EPC-9 amplifier (HEKA Elektronik, Lambrecht, Germany) controlled by PatchMaster software (HEKA Elektronik, Lambrecht, Germany). Patch-clamp electrodes were pulled and fire polished to 3.0±0.5 MΩ resistance for whole cell experiments, and 2.0±0.5 MΩ for pressure-clamp experiments, using a DMZ-Universal Puller (Zeitz Instruments, Munich, Germany). An AgCl wire was used as a reference electrode. Series resistance was electronically compensated. Whole-cell currents of mPTCs were acquired using repetitive 400 ms voltage ramps (at 2 sec intervals) from −120 mV to +120 mV, applied from a holding potential of 0 mV. Currents were sampled at 20 kHz and digitally filtered at 2.9 kHz. For statistical analysis, the currents were normalized to cell capacitance. No statistical difference was observed between the membrane capacitance of mPTCs *Trpv4*^+/+^ (25.0 ± 5.2 pF; n = 30) and *Trpv4*^−/-^ (27.2 ± 6.4 pF; n = 27). Solutions were applied to the cells via a home-made gravity-fed perfusion system, connected by a 5-way manifold, to a RC25 perfusion chamber (Warner Instruments, Hamden, CT, USA). For whole cell recordings, the standard extracellular solution had the following composition (in mM): 150 NaCl, 5 CsCl, 1 MgCl_2_, 1.8 CaCl_2_, 10 glucose, and 10 HEPES, buffered at pH 7.4 with NaOH. The osmolarity of this solution was 320±5 mOsm/kgH_2_O. For hypotonic cell swelling, we superfused mPTCs with a solution containing 105 mM NaCl, 5 mM CsCl, 1 mM MgCl_2_, 1.8 mM CaCl_2_, 10 mM glucose, 80 mannitol and 10 mM HEPES, buffered to pH 7.4 with NaOH (320±5 mOsm). Hypotonic cell swelling was induced by removing mannitol to lower the extracellular osmolarity to 240±5 mOsm. For hypotonic cell swelling in a ‘low-chloride’ containing extracellular solution, 105 mM NaCl was replaced by 105 mM Na-Aspartate. The pipette solution was composed of (in mM) 40 CsCl, 100 Cs-aspartate, 1 MgCl_2_, 10 HEPES, 4 Na_2_ATP, 10 EGTA, and 0.5 CaCl_2_, adjusted to pH 7.2 with CsOH (290±5 mOsm). For pressure-clamp recordings on mPTCs we used the following pipette solution (in mM), as described in (Peyronnet et al., 2013): NaCl 150, KCl 5, CaCl_2_ 2 and HEPES 10 (pH 7.4 with NaOH). The pipette solution also contained 10 mM TEA, 5 mM 4AP and 10 µM glibenclamide to inhibit K^+^ channels. The bath medium contained (in mM): KCl 155, EGTA 5, MgCl_2_ 3 and HEPES 10 (pH 7.2 with KOH), to zero the membrane potential. Membrane patches in cell-attached configuration were stimulated, at the holding potential of −80 mV, with 300 ms negative pressure pulses from 0 mmHg to −80 mmHg (10 mmHg steps, at 4 sec intervals), using a fast pressure-clamp device (ALA High Speed Pressure Clamp-1 System, ALAScientific). Data and statistical analysis was performed using Origin 8.0 (OriginLab Corporation, Northampton, MA).

### Albumin and dextran uptake assay

mPTCs were serum-starved overnight and incubated with 0.5 mg/mL FITC-labelled albumin (Sigma) or 1 mg/ml FITC-labelled dextran (Sigma) in DMEM/F12 medium for 30 min at 37°C. The cells were washed with cold PBS five times and lysed in PBS 0,1% Triton-X. The supernatant was used for fluorescence and protein assays. The intensity of FITC-fluorescence was measured by fluorophotometry. For the study of albumin endocytosis during cell stretching, cells were seeded on collagen I-coated flexible silicon chambers (ST-CH-10, B-Bridge) and placed in the stretching instrument (ST-140, B-Bridge). A uniaxial stretching of 20% was applied at a frequency of 0.3 Hz. At the end of the experiments, the cells were washed 5 times with ice cold PBS and the fluorescence was immediately assessed using a fluorescent microscope.

### AT2 Infusion Studies

A 14-days infusion of AT2 (Tocris) at a dose of 1 ng/g/min dissolved in sterile saline was performed using osmotic mini-pumps (Alzet model 2004; Alza Corp., Mountain View, CA) (Eckel et al., 2011). The pumps were implanted subcutaneously in the dorsal region under ketamine-xylazine anesthesia (100-10µg/g). After a recovery period of 4 days, 24-hour urine was collected at 4, 7 and 14 days post-implantation in metabolic cages.

### Anti-GMB serum treatment

To induce a progressive glomerulonephritis, 100 µl of anti-GBM serum (Sheep anti-Rat glomeruli serum, Probetex, Inc.) were administered to 6–8 week old mice by tail vein injection. Urine samples were collected on days 1, and 7 after antiserum injection. On day 7 the mice were sacrificed to collect plasma and kidney tissue.

### Urine analysis

After collection, 24-hour urine was centrifuged to remove debris, weighed and frozen at −80°C. Urine samples were analyzed for quantitative estimation of albumin (ab108792 ELISA kit, Abcam), glucose (Fuji Dri-Chem NX500 analyzer), phosphate (Phosphate Assay Kit, Abcam), creatinine (QuantiChromTM Creatinine Assay Kit, BioAssay Systems) and CC16 (Cloud Clone Corp.).

### Statistical analysis

All values are expressed as mean ± SEM. Paired t-test was used to assess statistical differences between two groups, ANOVA with Bonferroni’s test was used for multiple group comparisons. Statistical significance was indicated if P < 0.05.

## Acknowledgements

This work was supported by the Belgian Fund for Scientific Research (FNRS, grants CDR J0080.17 and EQP U.N011.17) and the Concerted Research Action from the General Direction of Scientific Research of the French Community of Belgium (ARC17/22-083) and the Swiss National Science Foundation (project grant 31003A-169850). FT is Research Director and FS Research Fellow of the FNRS.

## Author Contributions

RG, FS, OD and PG designed the study, performed experiments, interpreted data and wrote the paper. XY, OS and YA performed experiments and interpreted data. NT and FD critically revised the manuscript. All authors provided final approval for the version of the manuscript submitted for publication and agree to be accountable for the work.

